# Beyond Capture Efficiency: A Multidimensional Framework for Benchmarking Circulating Tumor Cell Isolation Technologies

**DOI:** 10.64898/2026.05.05.722894

**Authors:** C. Z. Valega Negrão, H. Hendrick, F. Ammar, R. Klotz, S. Dias, M. Yu

## Abstract

Metastasis remains the major cause of cancer-related mortality, and circulating tumor cells (CTCs) are both candidate liquid-biopsy biomarkers and plausible intermediates of metastatic dissemination. Because CTCs are extremely rare in peripheral blood, platform comparisons have often focused solely on recovery. That focus is insufficient for applications that depend on the quality of the recovered material, including single-cell profiling, short-term culture, and functional testing. Here, we compared four CTC isolation approaches: TellDx CTC System, Genesis System, RosetteSep™, and flow cytometry, using spike-in experiments in human blood. Capture efficiency was evaluated across all four platforms; purity was assessed for TellDx, Genesis, and RosetteSep™; and post-isolation GFP signal persistence in culture was assessed for TellDx and Genesis as an exploratory proxy for short-term post-isolation preservation. Under the conditions tested, TellDx showed the highest recovery (88.1% ± 3.7%), followed by Genesis (40.6% ± 12.1%), RosetteSep™ (36.5% ± 9.0%), and flow cytometry (7.6% ± 4.5%). TellDx also showed the highest purity score (3.76), whereas Genesis (2.25) and RosetteSep™ (2.09) did not differ substantially. In the short-term culture assay, TellDx-derived samples retained a higher normalized GFP signal than Genesis-derived samples at 48 h and 72 h. To synthesize these readouts, we propose the Recovery Performance Index (RPI), a composite score integrating recovery, purity, and post-isolation signal persistence. Within this experimental framework, TellDx achieved the highest RPI. These data support two conclusions. First, platform benchmarking for CTC workflows benefits from multidimensional evaluation rather than recovery alone. Second, under this spike-in model and within the specific workflows used here, TellDx performed best among the platforms tested. The principal contribution of this study is therefore the establishment of a practical benchmarking framework that can be expanded in future work using clinical samples, multiple CTC phenotypes, and orthogonal viability assays.

## Introduction

Metastasis accounts for most cancer-related deaths worldwide (1), and the bloodstream is a critical route for tumor dissemination (2). Circulating tumor cells (CTCs) are malignant cells shed from primary or metastatic lesions into the circulation (3), where only a small fraction persists long enough to contribute to secondary colonization (4). Despite their rarity, often only a few CTCs among billions of hematologic cells in a typical sample (5), CTCs have clear biological and clinical relevance: they provide access to tumor material through liquid biopsy (6), have prognostic value in several solid tumors (7,8), and may offer mechanistic insight into metastatic progression (9,10).

The isolation of CTCs remains technically demanding because these cells are vastly outnumbered by blood cells (5) and can exhibit marked phenotypic heterogeneity (11,12). Existing enrichment strategies rely on antigen expression, most notably targeting the epithelial cell adhesion molecule (EpCAM) as implemented in the CellSearch system (8), as well as on physical properties such as size and density, or on microfluidic-based separation approaches (13). In practice, however, platforms are often compared primarily based on capture efficiency (14). That emphasis is understandable, but it is incomplete. Recovery alone does not indicate whether the enriched population is sufficiently clean for downstream molecular analysis or whether the isolation procedure adequately preserves cells for short-term functional studies.

However, an important challenge in the field is not only to isolate CTCs but to do so while preserving their biological integrity for downstream functional studies (15). Beyond detection and enumeration, maintaining viable CTCs is essential for applications such as ex vivo culture, drug testing, and mechanistic analyses (15,16). Despite the development of multiple isolation platforms, systematic comparisons remain limited, and the immediate impact of different isolation strategies on CTC viability is still poorly understood (13). The lack of standardized evaluation frameworks further complicates the identification of approaches best suited for functional applications.

In this context, performance requirements extend beyond recovery alone. To address this gap, we define three operational dimensions: the number of candidate CTCs recovered, the extent of leukocyte carryover after enrichment, and the ability of recovered cells to tolerate the isolation workflow. A platform that maximizes recovery but introduces substantial contamination or imposes biological stress may be less suitable for downstream applications. Conversely, approaches that better preserve cell integrity, even with moderate recovery, may offer advantages depending on the intended use. We then integrated these parameters into a composite benchmarking metric, the Recovery Performance Index (RPI). We emphasize that this study was designed as a controlled comparative analysis in a spike-in model. It does not replace validation in clinical samples, but it provides a structured framework for comparing isolation workflows in a way that better reflects their suitability for downstream functional applications than recovery alone.

## Materials and Methods

### CTC culture

The CTC line BRx68 was derived from patients with luminal breast cancer, as previously reported (16). CTCs (female donors) were cultured in ultra-low attachment plates (Corning, Cat. CLS3471) using RPMI-1640 medium (Sigma-Aldrich) supplemented with EGF (20 ng/mL, Peprotech), bFGF (20 ng/mL, Peprotech), B27 (1×, Gibco, Cat. 17594-044), and antibiotic/antimycotic (1×, Sigma-Aldrich, Cat. 15240062), under hypoxic conditions (4% O_2_, 5% CO_2_).

### Spike-in model

To simulate CTCs in blood samples, BRx68 GFP-positive CTC was generated via lentiviral transduction of GFP-luciferase plasmid (9) cultured in vitro were spiked into whole human blood obtained from three healthy donors collected in EDTA-containing tubes (1.5 mg/mL). For workflows requiring peripheral blood mononuclear cell (PBMC) enrichment (e.g., RosetteSep and flow cytometry), PBMCs were isolated after spiking as described below. To enhance the brightness for the capture and purity assays, CTCs were labeled with CellTracker Green CMFDA (Invitrogen, Cat. C7025) according to the manufacturer’s instructions. Unless otherwise specified, 100 BRx68 CTCs per mL of blood were spiked into each sample. Each workflow was implemented according to its recommended operating conditions.

### TellDx CTC System (TellBio)

Whole blood samples spiked with CTCs were used to evaluate the TellDx CTC System platform (TellBio). Briefly, 1.2 mL of Dynabeads MyOne Streptavidin T1 (Invitrogen, Cat. 01-2222-41) were washed three times with 1.2 mL of 0.01% (v/v) Tween-20 in PBS (1×), followed by four washes with 1.2 mL of 0.1% (m/v) BSA in PBS. All washing steps were performed using a magnetic separator. After the final wash, beads were resuspended in 0.1% BSA in PBS and stored at 4 °C until use. Spiked blood samples were incubated with anti-CD45 (R&D Systems, clone 2D1), CD66b (AbD Serotec, clone 80H3), and CD16 (BD Biosciences, clone 3G8) antibodies at a ratio of 10:1:1, as recommended by the manufacturer, for 30 minutes at room temperature. Samples were then incubated with Dynabeads MyOne Streptavidin T1 for 20 minutes at room temperature. Each assay was conducted using 8 mL of human blood, with antibody and bead concentrations kept constant across experiments. Input volumes were defined according to each platform’s recommended operating conditions to ensure optimal performance. After incubation, samples were diluted 1:1 (v/v) with 0.2% (m/v) Pluronic F68 in PBS (pH 7.4) and processed using the TellDx CTC System. Collected samples were centrifuged at 300 × g for 5 minutes at 4 °C, and pellets were resuspended in 100 µL of PBS.

### Genesis System, Celselect Slides Technology (Bio-Rad)

For this methodology, 4 mL of human blood spiked with BRx68 CTCs was used, as recommended by the manufacturer. The Bio-Rad Enrichment Kit 1.0 (Bio-Rad, Cat. CEL80110) was used following the manufacturer’s instructions. The microfluidic workflow was developed in collaboration with Bio-Rad Laboratories. After processing, the collected samples were centrifuged at 300 × g for 5 minutes at 4 °C, and the resulting pellet was resuspended in 100 µL of PBS.

### RosetteSep™ Human CD45 Depletion Cocktail

A total of 4 mL of whole blood containing 400 spiked CTCs was processed using the RosetteSep Human CD45 Depletion Cocktail (StemCell Technologies, Cat. 15162). Briefly, 50 µL/mL of RosetteSep™ cocktail was added, and the mixture was incubated at room temperature for 20 minutes. Samples were diluted 1:1 (v/v) with PBS supplemented with fetal bovine serum (FBS), then density-separated using 4 mL of Lymphoprep (StemCell Technologies, Cat. 18060) and centrifuged at 1200 × g for 20 minutes at 4 °C. The PBMC layer containing CTCs was collected, washed twice with PBS supplemented with 2% FBS, and centrifuged at 300 × g for 10 minutes per wash. Final pellets were resuspended in 100 µL of PBS.

### Flow cytometry

Flow cytometry was included as a sorting-based workflow following density gradient separation and PBMC isolation. PBMCs were isolated from spiked blood samples using Lymphoprep as described above. After centrifugation, cells were blocked with BD Human Fc Block (BD Pharmingen™, Cat. 564219) and stained with CD45-APC (BioLegend, clone 2D1, Cat. 368512) for 15 minutes at 4 °C. Samples were washed twice with FACS buffer (PBS supplemented with 2% FBS, 0.02% EDTA, and 0.01% sodium azide; filtered through a 0.22 µm membrane). Cells were then stained with 7-AAD (BioLegend, Cat. 640922) to exclude non-viable cells. Data acquisition was performed using a Cytek Aurora Spectral Cytometer, and analysis was conducted using FlowJo (v10.10). The gating strategy included: (1) singlet selection (FSC vs. SSC), (2) exclusion of non-viable cells (7-AAD), and (3) identification of GFP^+^ CD45^−^ cells corresponding to CTCs.

### Capture efficiency

Capture efficiency was calculated as the proportion of recovered CTCs relative to the total number of spiked cells (100 CTCs/mL). CTCs were identified by microscopy using CellTracker Green CMFDA fluorescence, and the entire recovered volume was analyzed for each method. All experiments were performed in three independent biological replicates.

### Purity assessment

Purity was defined as the fraction of CTCs relative to total nucleated cells after enrichment (Equation 1).

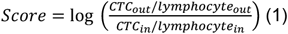

Lymphocyte counts prior to enrichment (lymphocytein) were determined using erythrocyte lysis with 3% acetic acid containing methylene blue (StemCell Technologies, Cat. 07060). After enrichment, CTC_out_ was determined based on fluorescence-positive cells, while fluorescence-negative cells were operationally classified as non-CTC contaminating cells (predominantly leukocytes, lymphocyte_out_). CTC_in_ and lymphocyte_in_ correspond to pre-enrichment counts, while CTC_out_ and lymphocyte_out_ correspond to post-enrichment counts.

### Post-isolation preservation assay

To evaluate short-term preservation after isolation, 1 × 10^5^ BRx68 GFP-positive CTCs were spiked into each sample according to the respective workflows, ensuring sufficient fluorescence signal for reliable detection. This input level was selected based on the fluorescence readout’s sensitivity limits. Two capture methodologies were evaluated: the Genesis System and the TellDx CTC System. Following isolation, recovered CTCs were transferred to 96-well plates and cultured as described above. GFP fluorescence intensity was measured at 24 h, 48 h, and 72 h using the Odyssey M Imaging System (LI-COR). Post-isolation survival was estimated by normalizing fluorescence signals at 48 h and 72 h relative to the signal measured at 24 h. Notably, GFP signal was used directly without additional fluorescent labeling. GFP signal was used as a proxy for cell persistence, assuming stable expression in viable cells over the analyzed time frame. These measurements should be interpreted as an exploratory proxy for short-term post-isolation persistence in culture rather than as a definitive assessment of cell viability.

### Recovery Performance Index (RPI)

To enable a multidimensional comparison of CTC isolation technologies, we developed a composite metric termed the Recovery Performance Index (RPI), integrating capture efficiency, sample purity, and post-isolation preservation. To combine these variables into a single comparative framework, a simplified application-driven multi-criteria decision analysis (MCDA) approach was applied.

Capture efficiency was expressed directly as a proportion (0–1 scale). Because purity and post-isolation preservation were measured on different scales, these parameters were normalized to the highest value observed among the evaluated platforms. The weighting scheme was set in advance to reflect each parameter’s importance for functional applications, not to produce a mathematically optimized model. Capture efficiency was assigned the highest weight, as it determines the number of CTCs available for downstream analyses. Post-isolation preservation was assigned to be the second-highest weight, reflecting its relevance for functional assays and short-term culture. Sample purity was also considered, given its importance in minimizing background signals in molecular analyses, although it was assigned a slightly lower weight.

The RPI was calculated as follows:

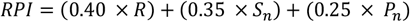

where *R* represents capture efficiency, *S*_*n*_normalized post-isolation preservation, and *P*_*n*_ normalized purity. RosetteSep™ and flow cytometry were excluded from the composite calculation because post-isolation preservation was not evaluated for these workflows. Accordingly, the RPI is presented as a proof-of-concept benchmarking tool rather than a definitive universal metric, and the weighting scheme can be adapted depending on the intended application.

### Statistical analysis

Data are presented as mean ± standard deviation. Cross-platform comparisons for capture efficiency and purity were performed using one-way ANOVA followed by Tukey’s post hoc test. The short-term post-isolation preservation assay was conducted as an exploratory study to compare two distinct workflows across sequential time points. Welch’s t-test was employed to evaluate between-group differences at each interval. All experiments were conducted in three independent biological replicates.

## Results

### Comparative evaluation of capture efficiency

We first compared the ability of four workflows to recover spiked BRx68 cells from human blood. Under the conditions tested, TellDx showed the highest capture efficiency (88.1% ± 3.7%), followed by Genesis (40.6% ± 12.1%), RosetteSep™ (36.5% ± 9.0%), and flow cytometry (7.6% ± 4.5%) (Figure 1). These differences indicate that, within this spike-in model, platform choice has a major effect on the fraction of input cells recovered after processing.

**Figure 1.**
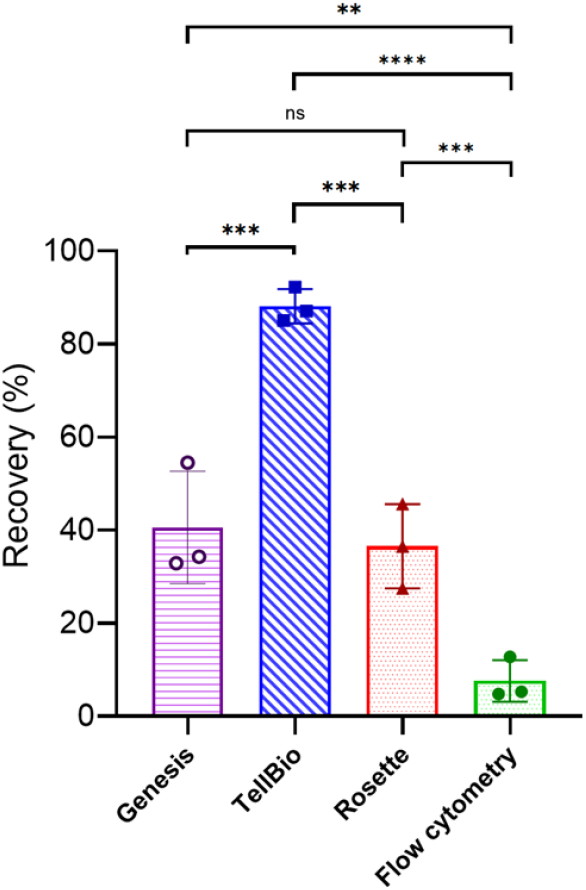
Capture efficiency of CTCs in human blood using four isolation workflows: instrument-based platforms (Genesis System and TellDx CTC System) and non-instrumented approaches (RosetteSep™ and flow cytometry). Data are presented as mean ± standard deviation (SD) from three independent biological replicates. Statistical analysis was performed using one-way ANOVA followed by Tukey’s multiple comparisons test. Significance levels are indicated as **p < 0.01, ***p < 0.001, ****p < 0.0001; ns, not significant.

The magnitude of the TellDx advantage in this dataset suggests that the combined use of antibody-functionalized beads and microfluidic processing may have contributed to the higher recovery observed under the tested conditions. At the same time, these values should be interpreted carefully. Recovery reflects the full workflow used for each platform, including sample handling, enrichment, transfer steps, and the specific operating conditions adopted here. The present data, therefore, supports a relative comparison within this study, not a platform-independent ranking across all possible implementations or sample types.

Controlled spike-in models using defined CTC concentrations are widely employed in methodological studies to enable reproducible quantification of recovery efficiency (17–21). Accordingly, the concentrations used here are higher than those typically observed in clinical samples and should be interpreted as a standardized benchmarking condition rather than a direct surrogate for clinical sensitivity.

While previous studies have compared selected CTC isolation strategies (20,21), including approaches based on antigen expression and biophysical properties, these comparisons have typically been limited to specific subsets of technologies or experimental conditions. For example, Drucker et al. evaluated immunomagnetic and size-based enrichment methods and reported differences in recovery performance across platforms. More recent studies have expanded these comparisons to additional technologies (21); however, direct benchmarking of multiple workflows under matched experimental conditions remains heterogeneous across the field. In this context, our study provides a direct head-to-head comparison of four distinct CTC isolation approaches within a unified experimental framework.

### Purity of Enriched samples

We next examined whether the platforms differed in the extent of non-CTC blood-cell carryover after enrichment. Among the three workflows evaluated for this endpoint, TellDx yielded the highest purity score (3.76), whereas Genesis (2.25) and RosetteSep™ (2.09) were lower and not significantly different from each other (Figure 2). These findings reinforce that recovery alone does not capture the full analytical value of an isolation workflow.

**Figure 2.**
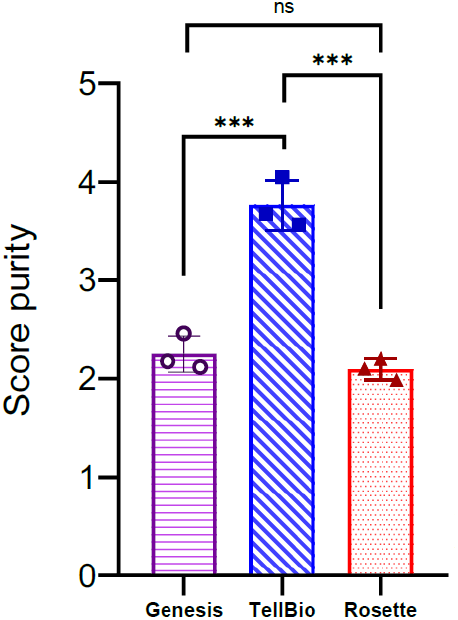
Purity of circulating tumor cell (CTC) enrichment across three isolation workflows: Genesis System, TellDx CTC System, and RosetteSep™. Purity is expressed as a score reflecting the ratio of recovered CTCs to residual leukocytes after enrichment. Data are presented as mean ± standard deviation (SD) from three independent biological replicates. Statistical analysis was performed using one-way ANOVA followed by Tukey’s multiple comparisons test. Significance levels are indicated as ***p < 0.001; ns, not significant

A plausible explanation for the improved TellDx purity is that the workflow combines target cell enrichment with more effective antibody-mediated depletion of background blood cells. By contrast, Genesis and RosetteSep™ rely on different separation principles, primarily microfluidic and density-based approaches, respectively, which may result in higher leukocyte carryover under the conditions tested. This distinction is particularly relevant for downstream molecular analyses, where contaminating leukocytes can dilute tumor-derived signals and complicate data interpretation.

### Short-term post-isolation preservation

TellDx and Genesis, which performed best in recovery and purity tests, were chosen for short-term post-isolation analysis. GFP fluorescence intensity was monitored at 24 h, 48 h, and 72 h after isolation as a proxy for persistence of the recovered cells in culture (Figure 3).

**Figure 3.**
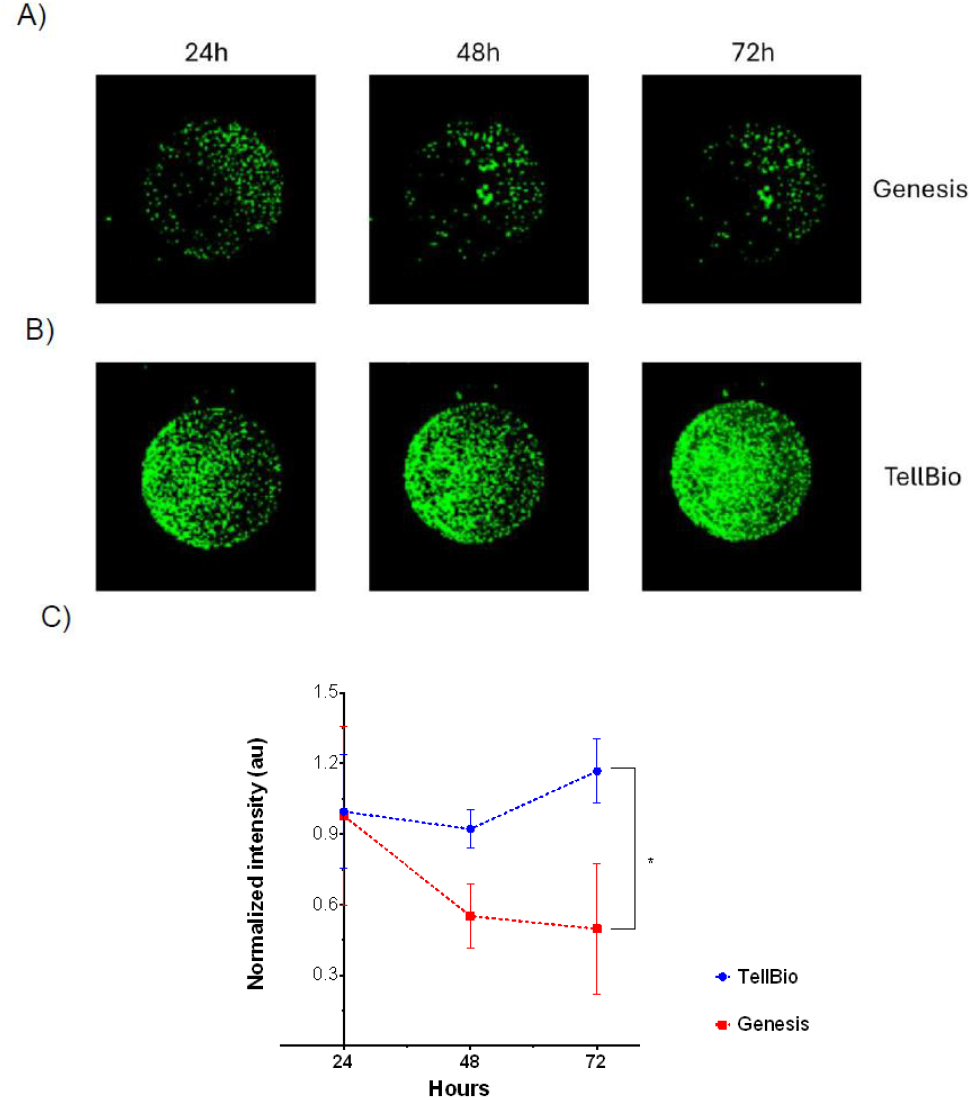
Short-term post-isolation preservation of circulating tumor cells (CTCs) recovered using TellDx and Genesis workflows. GFP fluorescence intensity was monitored at 24 h, 48 h, and 72 h after isolation and normalized to the 24 h time point. Data are presented as mean ± standard deviation (SD) from three independent biological replicates. Statistical comparisons at each time point were performed using Welch’s t-test. *p < 0.05.

Samples isolated with TellDx retained a higher normalized GFP signal over time, whereas those isolated with Genesis showed a progressive decline. At 48 h, the normalized signal was 0.92 ± 0.08 for TellDx and 0.55 ± 0.14 for Genesis (p = 0.033). At 72 h, the corresponding values were 1.17 ± 0.14 and 0.50 ± 0.27, respectively (p = 0.038). Within the limits of this assay, these results suggest that the TellDx workflow either imposed less stress on the recovered cells or better-preserved conditions compatible with short-term persistence in culture.

This endpoint should nevertheless be interpreted conservatively. The assay relies on GFP signal as a proxy and does not constitute a direct orthogonal measurement of viability, apoptosis, proliferation, or clonogenic potential. Accordingly, the results support a relative difference in post-isolation preservation between the two workflows but do not establish superior biological viability in a broader sense.

### Multidimensional benchmarking and the Recovery Performance Index

To integrate the three measured dimensions recovery, purity, and short-term post-isolation preservation, we performed a multidimensional benchmarking analysis using a radar plot and a composite metric, the Recovery Performance Index (RPI) (Figure 4A). The RPI was calculated using a simplified application-driven multi-criteria decision analysis framework, in which normalized values for each parameter were combined using a predefined weighting scheme reflecting their relevance for downstream applications.

**Figure 4.**
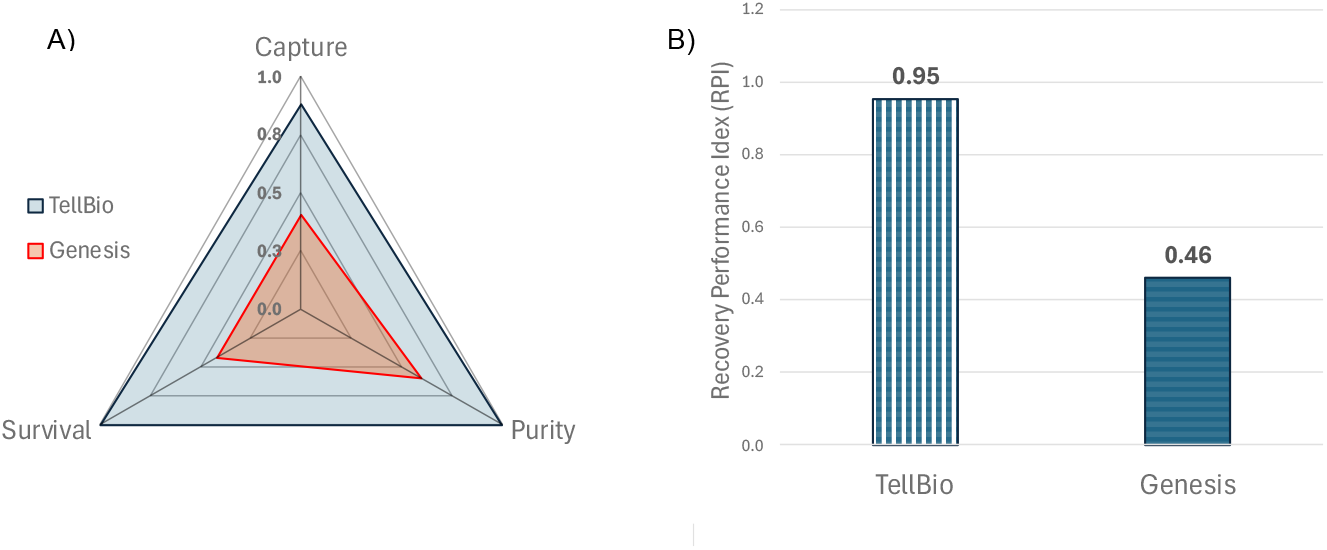
Multidimensional benchmarking of CTC isolation workflows. (A) Radar plot illustrating the relative performance of the evaluated platforms across capture efficiency, sample purity, and short-term post-isolation preservation. Values are normalized on a 0–1 scale. (B) Recovery Performance Index (RPI) derived from a weighted multi-criteria framework integrating these three parameters. The RPI is presented as a proof-of-concept composite metric for comparative evaluation of workflows.

Across the evaluated dimensions, TellDx consistently ranked highest, with the highest normalized values for all three parameters, whereas Genesis demonstrated intermediate performance. Using this framework, TellDx achieved an RPI of 0.95, compared with 0.46 for Genesis (Figure 4B). We emphasize that this score should be interpreted as a proof-of-concept composite metric. Its primary value lies in illustrating that benchmarking of CTC isolation technologies can be multidimensional and application-driven, rather than based solely on recovery. Because the current implementation depends on normalization within the tested set and excludes platforms for which all dimensions were not measured, the RPI should not be interpreted as a definitive universal ranking tool.

## Discussion and Future Perspectives

This study was designed to test a simple but important premise: benchmarking CTC isolation workflows based solely on recovery is insufficient for applications that depend on both the analytical quality and the biological state of the recovered cells. The data support that premise. Within the spike-in model used here, with assay-specific input conditions to ensure adequate detection sensitivity, platform performance was not defined solely by capture efficiency; purity and short-term post-isolation preservation further differentiated the workflows and influenced the practical interpretation of the results.

Under the specific workflows and experimental conditions applied here, TellDx showed the strongest overall performance across the measured dimensions. This observation is meaningful but should be interpreted in context. The present study does not establish an absolute hierarchy of CTC isolation technologies. Rather, it shows that, in a controlled spike-in setting using BRx68 cells, TellDx produced the most favorable combined profile among the tested workflows. This distinction is relevant because CTC isolation performance depends on multiple factors beyond the underlying separation principle, including cell phenotype, antigen expression, deformability, sample composition, and workflow implementation.

The RPI proposed here provides a structured framework to formalize this multidimensional view of performance. At the same time, the metric has limitations that should be made explicit. The weighting scheme is rational but investigator-defined; normalization depends on the specific set of platforms included, and only two workflows could be incorporated into the composite due to the availability of preservation data. Accordingly, the RPI should be viewed as a practical framework for structured comparison rather than as a finalized or universally applicable standard.

Some final considerations help contextualize these findings. First, the study relied on a single patient-derived CTC model, which, although biologically relevant, does not capture the full diversity of clinical CTC populations across different cancer types. Second, the spike-in design employed defined cell inputs exceeding typical CTC abundance in patient samples; while appropriate for controlled benchmarking, this approach does not fully reproduce the heterogeneity and scarcity of clinical specimens. In addition, post-isolation preservation was assessed using GFP fluorescence as an indirect readout, which cannot distinguish between survival, metabolic activity, or proliferation. Finally, the evaluated workflows differ not only in their isolation principles but also in sample handling and operational conditions, meaning that the results reflect the complete workflows as implemented.

Despite these limitations, the study provides a practical framework for comparing CTC isolation technologies under controlled conditions. The results highlight that recovery, purity, and preservation represent distinct and complementary dimensions of performance that may influence the suitability of a given platform for different experimental objectives. In this context, the primary contribution of this work is not the identification of a single “best” platform, but the demonstration that meaningful benchmarking requires a multidimensional perspective.

Future work should extend this framework to clinically relevant samples with lower CTC abundance, incorporate multiple CTC phenotypes and cluster-containing preparations, and include orthogonal assays to more directly assess cell viability and biological function. In addition, evaluating how alternative weighting schemes influence composite scores may further refine the use of multidimensional benchmarking in this field. Such studies will help establish more robust and context-dependent criteria for selecting CTC isolation technologies in both research and clinical settings.

## Conclusion

CTC isolation technologies should not be evaluated based on capture efficiency alone when the intended applications depend on both sample purity and preservation of recovered cells. In this controlled spike-in study, TellDx showed the strongest overall profile among the tested workflows, with higher recovery, higher purity, and improved short-term post-isolation GFP signal persistence compared with the primary comparator evaluated across all three dimensions. The broader contribution of this work, however, is the demonstration that multidimensional benchmarking is both feasible and more informative than single-metric comparisons. In this context, the Recovery Performance Index provides a practical proof-of-concept framework that can be further refined and validated in future studies using clinical samples and orthogonal functional readouts.

## Author contributions

C. Z. Valega Negrão contributed to conceptualization, experimental design, experimental execution, data analysis, manuscript drafting, and revision. H. Hendrick contributed to experiments, including TellDx and flow cytometry workflows, and to manuscript revision. F. Ammar assisted with flow cytometry experiments and manuscript revision. R. Klotz contributed to experimental design, CTC culture work, and manuscript revision. S. Dias and M. Yu supervised the study and critically revised the manuscript.

## Acknowledgements

Financial support was received from the National Council for Scientific and Technological Development (CNPq) (C.Z. Valega Negrão) and by funds through the Maryland Department of Health’s Cigarette Restitution Fund Program (CH-649-CRF). We thank the UMGCCC Flow Cytometry Shared Resource for their technical assistance. The authors have no other relevant affiliations or financial involvement with any organization or entity with a financial interest in or financial conflict with the subject matter or materials discussed in the manuscript apart from those disclosed.

## References

1. Breast cancer [Internet]. [cited 2023 May 8]. Available from: https://www.who.int/news-room/fact-sheets/detail/breast-cancer

2. Carty NJ, Foggitt A, Hamilton CR, Royle GT, Taylor I. Patterns of clinical metastasis in breast cancer: an analysis of 100 patients. European Journal of Surgical Oncology (EJSO). 1995 Dec 1;21(6):607–8.

3. TR. A. A case of cancer in which cells similar to those in the tumours were seen in the blood after death. Aust Med J [Internet]. 1869 [cited 2023 May 8];14:146-. Available from: https://cir.nii.ac.jp/crid/1573105976092422016

4. Fidler IJ. Metastasis: Quantitative Analysis of Distribution and Fate of Tumor Emboli Labeled With 125I-5-Iodo-2′-deoxyuridine. JNCI: Journal of the National Cancer Institute [Internet]. 1970 Oct 1 [cited 2023 May 8];45(4):773–82. Available from: https://academic.oup.com/jnci/article/45/4/773/982007

5. Ring A, Nguyen-Sträuli BD, Wicki A, Aceto N. Biology, vulnerabilities and clinical applications of circulating tumour cells. Nature Reviews Cancer 2022 23:2 [Internet]. 2022 Dec 9 [cited 2026 Mar 15];23(2):95–111. Available from: https://www.nature.com/articles/s41568-022-00536-4

6. Gu X, Wei S, Lv X. Circulating tumor cells: from new biological insights to clinical practice. Signal Transduction and Targeted Therapy 2024 9:1 [Internet]. 2024 Sep 2 [cited 2026 Mar 15];9(1):226-. Available from: https://www.nature.com/articles/s41392-024-01938-6

7. Lin D, Shen L, Luo M, et al. Circulating tumor cells: biology and clinical significance. Signal Transduction and Targeted Therapy 2021 6:1 [Internet]. 2021 Nov 22 [cited 2023 May 9];6(1):1–24. Available from: https://www.nature.com/articles/s41392-021-00817-8

8. Janni WJ, Rack B, Terstappen LWMM, et al. Pooled analysis of the prognostic relevance of circulating tumor cells in primary breast cancer. Clinical Cancer Research [Internet]. 2016 May 15 [cited 2023 May 8];22(10):2583–93. Available from: https://aacrjournals.org/clincancerres/article/22/10/2583/122011/Pooled-Analysis-of-the-Prognostic-Relevance-of

9. Klotz R, Thomas A, Teng T, et al. Circulating tumor cells exhibit metastatic tropism and reveal brain metastasis drivers. Cancer Discov [Internet]. 2020 Jan 1 [cited 2025 Apr 29];10(1):86–103. Available from: https://pubmed.ncbi.nlm.nih.gov/31601552/

10. Pantel K, Alix-Panabières C. Liquid biopsy and minimal residual disease — latest advances and implications for cure. Nature Reviews Clinical Oncology 2019 16:7 [Internet]. 2019 Feb 22 [cited 2026 Apr 11];16(7):409–24. Available from: https://www.nature.com/articles/s41571-019-0187-3

11. Yu M, Bardia A, Wittner BS, et al. Circulating breast tumor cells exhibit dynamic changes in epithelial and mesenchymal composition. Science [Internet]. 2013 Feb 1 [cited 2024 Apr 30];339(6119):580–4. Available from: https://pubmed.ncbi.nlm.nih.gov/23372014/

12. Topa J, Richert J, Stokowy T, et al. Characterizing epithelial-mesenchymal transition-linked heterogeneity in breast cancer circulating tumor cells at a single-cell level. Mol Oncol [Internet]. 2025 Dec 1 [cited 2026 Apr 11];19(12):3685–705. Available from: /doi/pdf/10.1002/1878-0261.70132

13. Agarwal A, Balic M, El-Ashry D, Cote RJ. Circulating Tumor Cells: Strategies for Capture, Analyses, and Propagation. Cancer J [Internet]. 2018 Mar 1 [cited 2024 Apr 30];24(2):70. Available from: /pmc/articles/PMC5880323/

14. Eslami-S Z, Cortés-Hernández LE, Thomas F, Pantel K, Alix-Panabières C. Functional analysis of circulating tumour cells: the KEY to understand the biology of the metastatic cascade. Br J Cancer [Internet]. 2022 [cited 2026 Apr 11];127(5). Available from: https://pubmed.ncbi.nlm.nih.gov/35484215/

15. Khoo BL, Grenci G, Lim YB, Lee SC, Han J, Lim CT. Expansion of patient-derived circulating tumor cells from liquid biopsies using a CTC microfluidic culture device. Nat Protoc [Internet]. 2018 Jan 1 [cited 2026 Apr 11];13(1):34–58. Available from: https://pubmed.ncbi.nlm.nih.gov/29215634/

16. Yu M, Bardia A, Aceto N, et al. Ex vivo culture of circulating breast tumor cells for individualized testing of drug susceptibility. Science [Internet]. 2014 Jul 7 [cited 2024 Apr 30];345(6193):216. Available from: /pmc/articles/PMC4358808/

17. Fachin F, Spuhler P, Martel-Foley JM, et al. Monolithic Chip for High-throughput Blood Cell Depletion to Sort Rare Circulating Tumor Cells. Scientific Reports 2017 7:1 [Internet]. 2017 Sep 7 [cited 2025 Apr 29];7(1):1–11. Available from: https://www.nature.com/articles/s41598-017-11119-x

18. Riahi R, Gogoi P, Sepehri S, et al. A novel microchannel-based device to capture and analyze circulating tumor cells (CTCs) of breast cancer. Int J Oncol [Internet]. 2014 [cited 2026 Mar 15];44(6):1870–8. Available from: https://pubmed.ncbi.nlm.nih.gov/24676558/

19. Xu L, Mao X, Imrali A, et al. Optimization and Evaluation of a Novel Size Based Circulating Tumor Cell Isolation System. PLoS One [Internet]. 2015 Sep 23 [cited 2026 Mar 15];10(9). Available from: https://pubmed.ncbi.nlm.nih.gov/26397728/

20. M Saini V, Oner E, Ward MP, et al. A comparative study of circulating tumor cell isolation and enumeration technologies in lung cancer. Mol Oncol [Internet]. 2025 Jul 1 [cited 2026 Apr 11];19(7):2014–37. Available from: /doi/pdf/10.1002/1878-0261.13705

21. Drucker A, Teh EM, Kostyleva R, Rayson D, Douglas S, Pinto DM. Comparative performance of different methods for circulating tumor cell enrichment in metastatic breast cancer patients. PLoS One [Internet]. 2020 Aug 1 [cited 2025 Apr 29];15(8):e0237308. Available from: https://pmc.ncbi.nlm.nih.gov/articles/PMC7425969/

